# Rosuvastatin-Eluting Gold Nanoparticle-Loaded Perivascular Implantable Wrap for Enhanced Arteriovenous Fistula Maturation in a Murine Model

**DOI:** 10.1101/2023.02.02.526859

**Authors:** Carleigh Klusman, Benjamin Martin, Joy Vanessa D. Perez, Allan John R. Barcena, Marvin R. Bernardino, Erin Marie D. San Valentin, Jossana A. Damasco, Huckie C. Del Mundo, Karem Court, Biana Godin, Natalie Fowlkes, Richard Bouchard, Jizhong Cheng, Steven Y. Huang, Marites P. Melancon

**Author notes:** ***Address for Correspondence***, Unit 1471, 1515 Holcombe Blvd., Houston, TX 77030. ***Author Contributions*** Conceptualization, C.K., B.M., S.Y.H., M.P.M; investigation, C.K., B.M., J.V.D.P., A.J.R.B., M.R.B, E.M.D.S.V. J.A.D, H.C.D.M, K.C., B.G., N.F., R.B., J.C., S.Y.H., M.P.M; data curation, C.K., B.M., J.V.D.P., A.J.R.B., M.R.B, E.M.D.S.V. J.A.D, H.C.D.M, K.C., B.G., N.F., R.B., J.C., S.Y.H., M.P.M; formal analysis, C.K., B.M., J.V.D.P., A.J.R.B., M.R.B, E.M.D.S.V. J.A.D, H.C.D.M, K.C., B.G., N.F., R.B., J.C., S.Y.H., M.P.M; writing—original draft, C.K., B.M.; writing—review and editing, C.K., B.M., J.V.D.P., A.J.R.B., M.R.B, E.M.D.S.V. J.A.D, H.C.D.M, K.C., B.G., N.F., R.B., J.C., S.Y.H., M.P.M; visualization, C.K., B.M., J.V.D.P., A.J.R.B., M.R.B, E.M.D.S.V. J.A.D, H.C.D.M, K.C., B.G., N.F., R.B., J.C., S.Y.H., M.P.M; supervision, M.P.M.; project administration, M.P.M.; funding acquisition, S.Y.H., M.P.M.; All authors have read and agreed to the published version of the manuscript. ***Disclosures*** There are no relationships with the industry that require disclosure here.

## Abstract

**Background:** Arteriovenous fistulas (AVFs) are a vital intervention for patients requiring hemodialysis, but they also contribute to overall mortality due to access malfunction. The most common cause of both AVF non-maturation and secondary failure is neointimal hyperplasia (NIH). Absorbable polycaprolactone (PCL) perivascular wraps can address these complications by incorporating drugs to attenuate NIH, such as rosuvastatin (ROSU), and metallic nanoparticles for visualization and device monitoring.

**Objectives:** This study aimed to assess the impacts of gold nanoparticle (AuNP) and ROSU-loaded perivascular wraps on vasculature NIH and AVF maturation and patency in a chronic kidney disease rat model.

**Methods:** Electrospun wraps containing combinations of PCL, AuNP, and ROSU were monitored for *in vitro* drug elution, nanoparticle release, tensile strength, and cell viability. Perivascular wraps were implanted in chronic kidney disease rats for *in vivo* ultrasound (US) and micro-computed tomography (mCT) imaging. AVF specimens were collected for histological analyses.

**Results:** No difference in cell line viability was observed in ROSU-containing grafts. *In vitro* release studies of ROSU and AuNPs correlated with decreasing radiopacity over time on *in vivo* mCT analysis. The mCT study also demonstrated increased radiopacity in AuNP-loaded wraps compared with PCL and control. The addition of ROSU demonstrated decreased US and histologic measurements of NIH.

**Conclusions:** The reduced NIH seen with ROSU-loading of perivascular wraps suggests a synergistic effect between mechanical support and anti-hyperplasia medication. Furthermore, the addition of AuNPs increased wrap radiopacity. Together, our results show that radiopaque, AuNP-, and ROSU-loaded PCL grafts induce AVF maturation and suppress NIH while facilitating optimal implanted device visualization.

## INTRODUCTION

As detailed in the United States Renal Data System’s 2022 Annual Data Report, 14.0% of all United States adults are living with chronic kidney disease, and end-stage renal disease prevalence has increased by 107% since 2000, with over 800,000 adults currently diagnosed. In 2020 alone, over 110,000 end-stage renal disease patients initiated hemodialysis treatment, for a cumulative total of almost 600,000 United States patients relying on life-saving hemodialysis therapy (1). With the prevalence of disease states requiring hemodialysis rows, reliable arteriovenous access has become increasingly crucial. However, vascular access complications continue to be a common source of morbidity and mortality in this population, with current data reporting a 40.9% one-year failure rate for primary unassisted arteriovenous fistulas (AVF). At 18 months after hemodialysis initiation, only 64.2% remain on hemodialysis therapy, with morality accounting for 24.9% of treatment termination (1).

The leading complication of AVFs impacting both maturation and post-maturation patency is vessel stenosis (2). While the pathophysiology of stenosis is not fully understood, it is believed to be primarily driven by post-operative abnormal proliferation of the innermost vessel wall, known as neointimal hyperplasia (NIH) (2,3). Elevated low-density lipoprotein cholesterol levels were found to be an independent risk factor for AVF patency failure (4), leading to studies investigating the impact of systemic statin therapy on AVF maintenance. Statins provide other known benefits, such as the regulation of endothelial cell function and inhibition of vascular inflammation and thrombosis in end-stage renal disease patients (5). Systemic statins have demonstrated mixed results on AVF failure rates, with possible significant attenuation of NIH by specific medications within the class (6,7). Moreover, patients may experience adverse effects, such as myopathy and hepatotoxicity, from high-dose systemic therapy. Localized perivascular delivery can allow the delivery of high-dose treatment on the vascular wall without significantly increasing the risk for systemic toxicity (8). However, data is lacking for the targeted delivery of statin medications to the AVF site.

Previous research assessing the use of bioabsorbable wrap material has demonstrated its utility for facilitating the delivery of a desirable structural and cellular profile to the newly anastomosed vasculature required for AVF physiology. For example, polycaprolactone (PCL) has been used in recent murine model studies to introduce mesenchymal stem cells to the AVF site via perivascular wraps implanted during anastomosis creation (8). Importantly, monitoring the effects of these mechanical and physiologic interventions remains a challenge with existing non-invasive imaging modalities. For this reason, a solution for ensuring vessel patency is a priority for successfully implementing this technology.

Radiopaque nanoparticles, including gold nanoparticles (AuNP), provide an accessible solution for the current limitations of AVF device surveillance (9,10). The safety and efficacy of AuNPs for intravascular graft monitoring have been demonstrated in a previous study aimed at monitoring the precise positioning and integrity of inferior vena cava filters using standard radiographic and computed tomography imaging modalities (10). The incorporation of radiopaque nanoparticles into existing, physiologically compatible polymers has proven beneficial for other commonly used vascular devices, as seen with the increased visibility of resorbable, polydioxanone polymer inferior vena cava filters when implanted with Au and bismuth NPs (11–14). For this study, the supplementation of bioabsorbable PCL wraps with AuNPs can offer the advantage of monitoring the effects of local mechanical and pharmacological therapy embedded within the vascular graft.

This study aims to determine the impact of a specific statin, rosuvastatin (ROSU), on the attenuation of NIH in a murine model after the surgical creation of AVF. Unlike prior studies, which evaluated systemic statin medications, this study assessed targeted ROSU delivered directly to the site of the AVF via elution from an implanted perivascular polymeric wrap. As AuNPs have been shown to enhance imaging techniques for various indications, this study also aims to evaluate the effects of embedded AuNPs on improved visibility and device monitoring of implanted AVF perivascular wrap.

## METHODS

### Materials

Methanol (ACS reagent grade, ≥99.5%), toluene (ACS reagent grade, ≥99.9%), gold (III) chloride trihydrate (HAuCl_4_·3H_2_O, ≥99.9%), tetrabutylammonium bromide (TBAB, ACS reagent grade, ≥98.0%), 1-octanethiol (≥98.5%), PCL (average Mn 80,000) and sodium borohydride (NaBH_4_, ≥99%) were obtained from Sigma-Aldrich (St. Louis, MO). ROSU was purchased from Acros (NJ). All chemicals were used without further purification unless otherwise noted.

### Fabrication of polymeric wraps

AuNPs were synthesized using the protocol established by Tian et al. (14) HAuCl_4_·3H_2_O in distilled water was reacted in the presence of TBAB, 1-octanethiol, and NaBH_4_. The resulting AuNP solution was resuspended in 8:3 toluene/MeOH at 0%, 10%, 20%, and 30% and mixed with 15% PCL. ROSU was incorporated into the mixture at 40 mg/mL. Using a Spraybase electrospinning system (Avectas, Maynooth, Ireland), each of the four solutions was electrospun at 1.0 mL/h and 15 kV.

### Characterization

JEM 1010 transmission electron microscopy (JEOL USA, Inc., Peabody, MA) was used to characterize both size and morphology of AuNPs. Using a Nova NanoSEM scanning electron microscope (Field Electron and Ion Company, Hillsboro, OR) with an EDAX energy dispersive spectroscopy system (Ametek, Berwyn, PA), additional parameters, including fiber diameter and pore size of wraps were obtained. Notably, the mean fiber diameters and mean pore sizes recorded via ImageJ (National Institutes of Health, Bethesda, MD) reflect randomly selected areas of the polymeric wrap. A combination of calculation (15) and liquid intrusion (16) methods were used to determine porosity. STA PT 1000 thermogravimetric analysis (Linseis, Selb, Germany) enabled the determination of melting temperature and glass transition temperature. Ultimate tensile strength was recorded using an MTESTQuattro materials testing system (ADMET, Norwood, MA).

### In vitro Drug and Nanoparticle Release

The polymeric wraps were incubated in phosphate-buffered saline (PBS) at 37°C with vigorous shaking for up to 12 weeks to simulate the hydrolytic degradation of the polymers. PBS was sampled and replenished after each time point. The amount of ROSU released at each time point was calculated from the absorption of the supernatant at 244 nm using a UV-vis spectrophotometer (Cary 60, Agilent, Santa Clara, CA), while the AuNP content was quantified by inductively coupled plasma - optical emission spectrometry (Varian 720-ES, Agilent, Santa Clara, CA).

### In vitro Cytotoxicity Assay

Immortalized human vascular endothelial cells (EC-RF24) and mouse vascular smooth muscle cells (MOVAS) were cultured in Dulbecco’s Modified Eagle Medium with 10% fetal bovine serum and 1% penicillin-streptomycin at 37°C in a 5% CO2 atmosphere. The gas-sterilized wraps were immersed in cell culture media for 24 hours to produce the treated media. EC-RF24 and MOVAS were seeded into 96-well plates at 1×10^4^ cells/well and incubated for 24 hours. After which, the cell culture media was replaced with treated media at increasing concentrations (i.e., 0, 25, 75, and 100%). After 24 hours, cell viability was measured using the alamarBlue™ assay. Fluorescence readings were taken at an excitation wavelength of 540 nm and emission wavelength of 590 nm using a Cytation5 microplate reader (Biotek Instruments, Winooski, VT).

### Animal Handling

Approval and oversite from the MD Anderson Cancer Center Institutional Animal Care and Use Committee were obtained for all experiments using rats. Up-to-date institutional licensing and training were required for any personnel handling the animals. Rats were appropriately prepared for all procedures, with sterilization of the surgical site using alternating chlorhexidine and alcohol in three repetitions. Sterile technique, including surgical drapes and sterile gloves and instruments, was upheld during invasive procedures. The periprocedural optical ointment was utilized, and animals were positioned in a standardized supine posture on the sterile field, which overlayed a heating blanket. Isoflurane anesthesia was utilized, and stimulation of intratarsal skin was applied to ensure adequate sedation before animal placement on the operating table. Buprenorphine was administered for pain control. Rats were euthanized via CO_2_ asphyxiation followed by thoracotomy.

### AVF Creation & Perivascular Attachment

Briefly, a two-step 5/6^th^ nephrectomy procedure from Wang et al. (17) was performed using Sprague-Dawley rats to model chronic kidney disease *in vivo*. Following a 4-week interval, AVFs were surgically created based on a modified procedure by Wong et al. (18). Specifically, end-to-side surgical anastomosis of the external jugular vein with the common carotid artery was performed. For analysis, the rats were divided into 4 groups—control AVF (i.e., no wrap), PCL, PCL-Au, and PCL-Au-ROSU. In all groups, excluding the control, a treatment-specific, gas-sterilized cylindrical wrap measuring 1 mm x 5 mm was incorporated into the AVF. The wrap in each group was fashioned around the outflow vein and secured using 10-0 nylon sutures.

### X-Ray and Micro-Computed Tomography

X-ray and micro-computed tomographic (mCT) studies of the wraps were collected prior to *in vivo* application. An MX-20 digital specimen radiography system (Faxitron Bioptics, Tucson, AZ) was used for radiographic visualization. A CT-eXplore Locus RS preclinical *in vivo* scanner (GE Medical Systems, London, ON, Canada) was used for *in vitro* and *in vivo* mCT imaging of the wraps expressed in Hounsfield units (HU).

### Ultrasonography

Pulsatile arterial waveforms in the external jugular vein of successfully created AVFs were visualized using the Vevo 2100-LAZR system (VisualSonics, Toronto, ON, Canada). At 2-, 4- and 8-week (W) time points, living rats were anesthetized and analyzed using B-mode ultrasound (US) with a 15-MHz probe. AVF outflow track wall thickness (WT) and luminal diameter (LD) were assessed, both with 6 measurements analyzed per scan.

### Histological Analysis

Pre-selected rats in each group were euthanized at W2, W4, and W8 time points. The outflow veins were harvested after perfusion at the site of perivascular wrap placement for formalin fixation and paraffin embedding. Blocks were sectioned into 4-μm sections, stained with hematoxylin and eosin, and subsequently imaged using an Aperio LV1 real-time digital pathology system (Leica Biosystems, Buffalo Grove, IL). ImageJ software was used to quantify each image’s luminal area (LA) and neointimal area (NIA).

### Statistical Analysis

Descriptive statistics were conducted with quantitative data and presented as means ± standard deviations. Mixed-effects models were used to compare US-derived WT, LD, and WT:LD ratio and histology-derived LA, NIA, and NIA:LA ratio between control, PCL, PCL-Au, and PCL-Au-ROSU treatments and time points. Mixed effects models were also used to compare both EC-RF24 and MOVAS in percent cell viability by fluorescence between control, PCL, PCL-Au, and PCL-Au-ROSU treatments. Ordinary one-way ANOVA was used to compare the percent cumulative 12W AuNP and ROSU release between PCL, PCL-Au, and PCL-Au-ROSU conditions. Tukey’s Multiple Comparison Test was used to adjust for multiple comparisons on all analyses. Statistical significance was defined as p <.05. All statistical analyses were conducted using GraphPad Prism version 9.0.0 (GraphPad, San Diego, CA)

## RESULTS

### Physiochemical Characteristics

AuNPs synthesized for incorporation in select wraps were found to have an average diameter of 3.93 ± 0.68 nm. These nanoparticles were incorporated successfully into PCL using the electrospinning method. The representative scanning electron microscope, photographic, x-ray, and mCT images of PCL with AuNP (PCL-Au) and AuNP and ROSU (PCL-Au-ROSU) polymeric wraps are shown in figure 2. Our results show that qualitatively in scanning electron microscope images, PCL-Au-ROSU fibers are more aligned than the fibers of PCL and PCL-Au. The addition of AuNP renders the previously white wrap a dark gray color, as seen in photographic images, with increased visibility on X-ray and m-CT imaging. The physicochemical properties of the polymeric wraps are summarized in table 1.

**Table 1.** Physico-chemical Properties of the Electrospun Wraps. A preliminary analysis was performed to compare the basic physiochemical properties of each polymer. A combination of transmission electron microscopy and energy dispersive spectroscopy was used to determine the diameter and pore size of the various polymers. The additional parameters collected are porosity, melting temperature, glass transition temperature, and maximum tensile stress. Analysis of each property showed no significant differences between polymeric compositions. The absence of substantial changes in these properties among wraps is useful for understanding the effect of each polymeric composition beyond its isolated physical and chemical features when introduced intravascularly. Abbreviations: Au, gold; PCL, polycaprolactone; ROSU, rosuvastatin; T_g_, glass transition temperature; T_m_, melting temperature.

| | Diameter<br>($\mu\text{m}$) | Pore size<br>($\mu\text{m}^2$) | % Porosity (%) | | $T_g$<br>($^{\circ}\text{C}$) | $T_m$<br>($^{\circ}\text{C}$) | Maximum<br>stress (MPa) |
| --- | --- | --- | --- | --- | --- | --- | --- |
|  |  |  | Calc. | Intrusion |  |  |  |
| **PCL** | $1.07 \pm 0.52$ | $7.97 \pm 11.4$ | $93.21 \pm 1.23$ | $78.6 \pm 8.06$ | -<br>69.61 | 52.53 | $9.17 \pm 0.31$ |
| **PCL-Au** | $1.10 \pm 0.49$ | $11.8 \pm 17.1$ | $92.77 \pm 2.61$ | $81.2 \pm 16.90$ | -<br>69.71 | 52.44 | $9.01 \pm 0.42$ |
| **PCL-Au-ROSU** | $0.99 \pm 0.49$ | $6.83 \pm 10.3$ | $92.27 \pm 1.50$ | $67.9 \pm 5.99$ | -<br>69.15 | 54.23 | $5.81 \pm 0.55$ |

**Figure 1.**
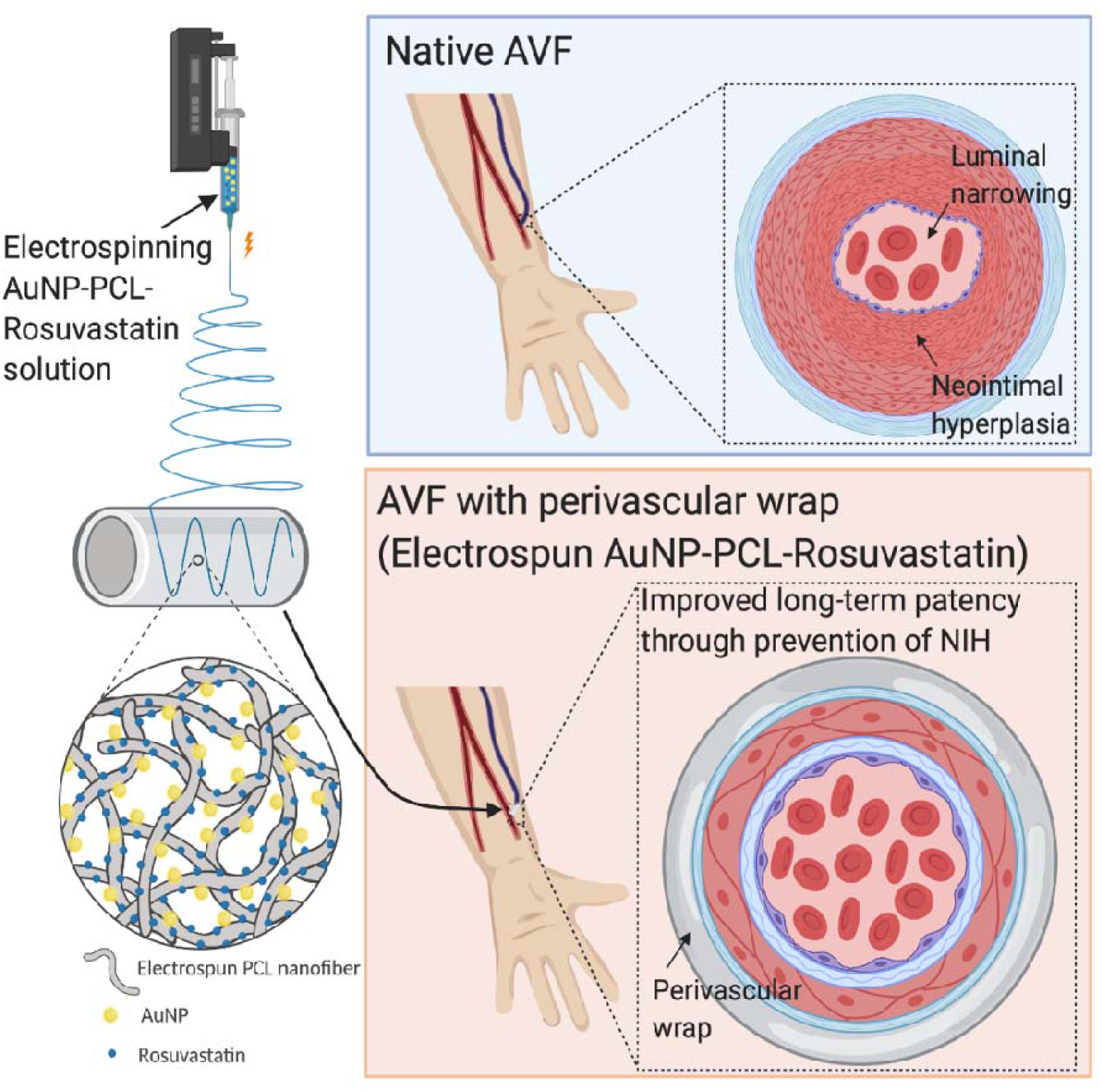
Central Illustration.

**Figure 2.**
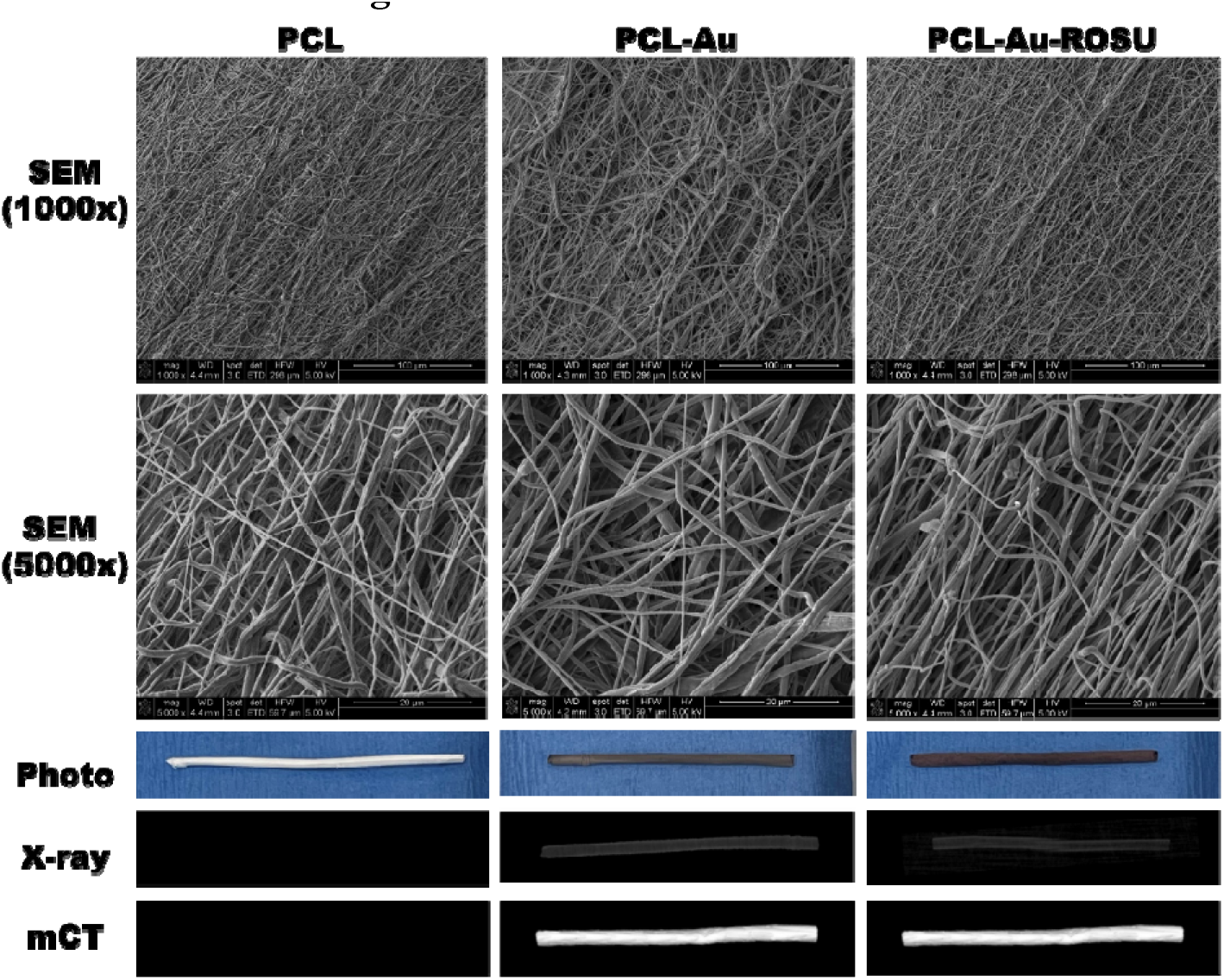
Physico-chemical Characterization and Imaging of the Elecrospun Wraps. Representative scanning electron microscope (1000x, bar=100 um; 5000x, bar=20 um), photographic, x-ray, and mCT images of the electrospun wraps are organized to compare each polymeric composition visually. Analysis demonstrates a greater alignment of fibers in the PCL-Au-ROSU wrap relative to PCL or PCL-Au. Incorporating gold nanoparticles results in a characteristic gray color in photographs and enhanced visualization on X-Ray and mCT imaging. Multimodal visual inspection is required for correlation with additional physiochemical properties of these polymers and intravascular performance when introduced as perivascular wraps to arteriovenous fistulas. Abbreviations: Au, gold; mCT, micro-computed tomographic; PCL, polycaprolactone; ROSU, rosuvastatin; SEM, scanning electron microscope.

Quantitatively, PCL-Au wraps had the largest average pore size out of all three wrap conditions (p≤0.013). PCL-Au-ROSU demonstrated the smallest average pore size but was not significantly different from bare PCL (p=0.613). Although intrusion porosity was similarly the highest in PCL-Au and lowest in PCL-Au-ROSU, no significant differences were found (p≥0.381). The three wraps also demonstrated no significant difference in fiber diameter (p≥0.513).

### In vitro ROSU and AuNP release from perivascular wraps and the impact of ROSU-loading on cell viability

*In vitro* % AuNP and % ROSU release rates are detailed in Supplementary tables 1 and 2 and plotted in figure 3, which compares substrate release over time across each polymeric wrap - PCL alone, PCL-Au, and PCL-Au-ROSU. *In vitro* % ROSU release demonstrated a significantly greater % cumulative release from W1-12 in the PCL-Au-ROSU group compared to the PCL and PCL-Au groups (p<0.0001). The ROSU release rate exhibited an initial burst during W1 that tapered over W2-12 for a total cumulative ROSU release of 40.7% at W12.

**Figure 3.**
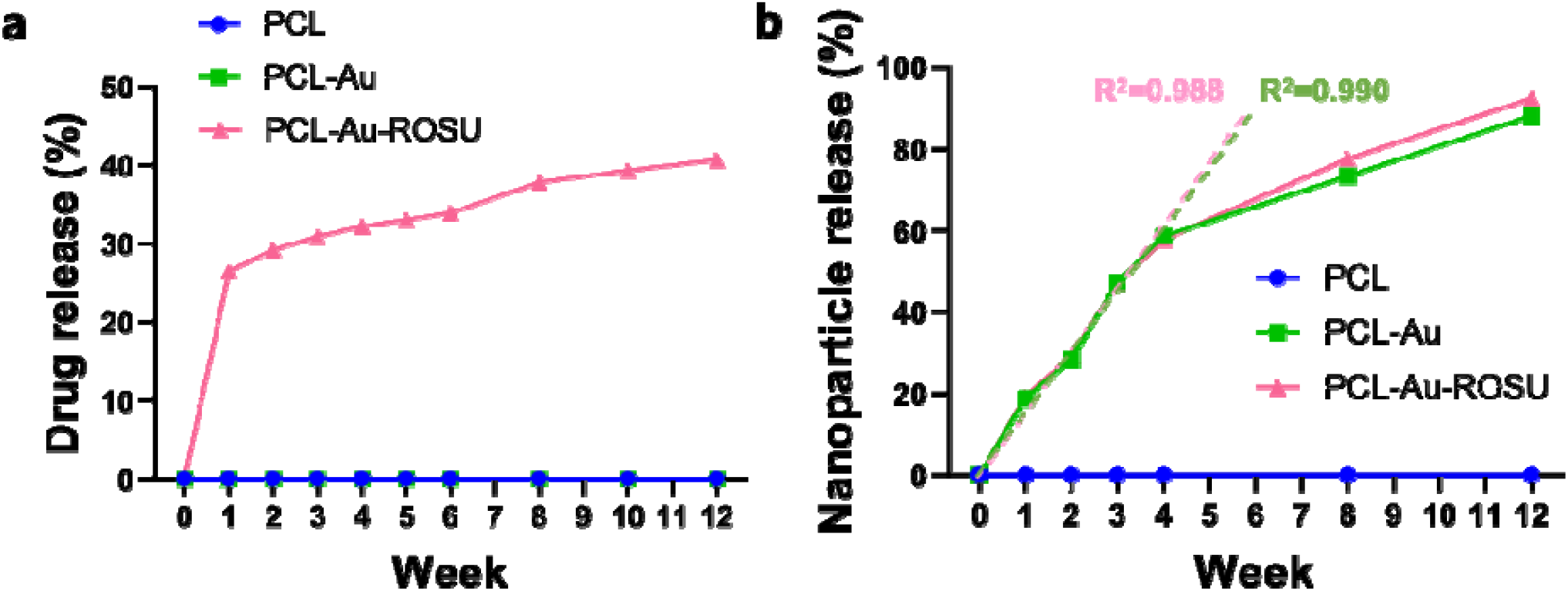
In Vitro Drug and Nanoparticle Release. PCL wraps were prepared under three loading conditions - bare PCL, PCL-Au, and PCL-Au-ROSU - and incubated in PBS to replicate in vivo device degradation over 12 weeks. Analysis of % cumulative (a) ROSU and (b) AuNP at various time points are depicted, with overlayed linear models, and R2 values included for PCL-Au and PCL-Au-ROSU AuNP release from weeks 1-4. ROSU demonstrated an initial release burst, followed by a gradual taper over weeks 1-12. AuNPs followed a linear release pattern from weeks 1-4, which suggests the potential for predictive models of implanted device degradation via clinical imaging. No difference in AuNP release rate was observed between PCL-Au and PCL-Au-ROSU, projecting no variation of in vivo radiopacity trends with the addition of ROSU to AuNP-loaded wraps. Abbreviations: PCL, polycaprolactone; Au, gold; ROSU, rosuvastatin; PBS, phosphate-buffered saline; AuNP, gold nanoparticle.

There was no difference in *in vitro* % cumulative AuNP release between PCL-Au and PCL-Au-ROSU (p=0.9952), both of which had greater AuNP release than PCL alone (p≤0.0125). Unlike ROSU, which had a 26.5% release burst at W1 followed by a gradual taper, average AuNP release was only 19.4% at W1 and then had a linear increase until W4 in both PCL-Au (R^2^=0.9901) and PCL-Au-ROSU (R^2^=0.9992). AuNP release slowly plateaued from W4 to W12 with a cumulative % AuNP release of 88.2% for PCL-Au and 92.4% for PCL-Au-ROSU.

Figure 4 demonstrates *in vitro* EC-RF24, and MOVAS fluorescence for the PCL-Au-ROSU compared to control well, PCL, and PCL-Au when stratified by percent of media treated. No significant difference in fluorescence between PCL, PCL-Au, and Au-PCL+ROSU treatment conditions was seen in EC-RF24 or MOVAS cell lines at any % media treated (p>0.05).

**Figure 4.**
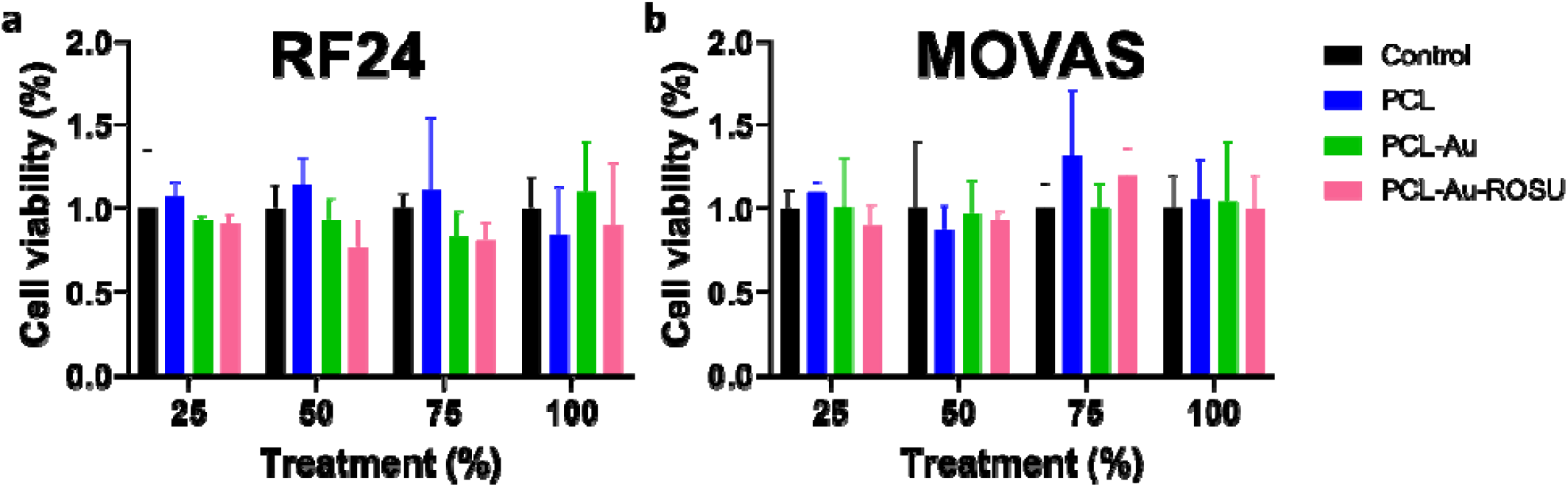
In Vitro Cell Viability Assay. (a) EC-RF24 endothelial cells and (b) MOVAS smooth muscle cells after 48 hours of incubation. Cell viability of PCL-Au-ROSU wraps, stratified by % of media treated, were compared to control wells, PCL wraps, and PCL-Au wraps to determine if the presence of ROSU impacted EC-RF24 or MOVAS lines. There was no difference in cell viability of EC-RF24 or MOVAS lines between PCL-Au-ROSU and any comparison groups at all concentrations analyzed, suggesting that ROSU-addition is not cytotoxic to these lines. Abbreviations: Au, gold; PCL, polycaprolactone; ROSU, rosuvastatin; EC-RF24, immortalized human vascular endothelial cells; MOVAS, mouse vascular smooth muscle cells

### AuNP-loaded grafts are associated with increased radiopacity on mCT of CKD rats

Figure 5(a) depicts *in vivo* representative mCT images of control, PCL, PCL-Au, and PCL-Au-ROSU at W2, W4, and W8. Average radiopacities in HU are detailed in table 2. Polymeric wraps loaded with AuNPs, such as PCL-Au and PCL-Au-ROSU, were found to be more radiopaque than control and PCL at all time points, including W2 (p≤0.0011), W4 (p<0.0001) and W8 (p<0.0001). In addition, PCL-Au-ROSU demonstrated a significantly higher HU than PCL-Au (p=0.0003) at W8. There was no difference between the AuNP-loaded wraps at W2 or W4 (p≥0.4869). PCL was found to be more radiopaque than the control at W2 (p=0.0028), but no difference between the two groups was seen at W4 or W8 (p≥0.0817). Figure 5(b) shows that the average radiopacities of AuNP-containing wraps PCL-Au and PCL-Au-ROSU decreased from W2 to W8. However, they remained significantly more visible than without AuNP throughout the study. This decrease in average HU was found to be linear to W4 for PCL-Au (R^2^=0.991) and PCL-Au-ROSU (R^2^=0.999) as shown in Supplementary Figure 1.

**Figure 5.**
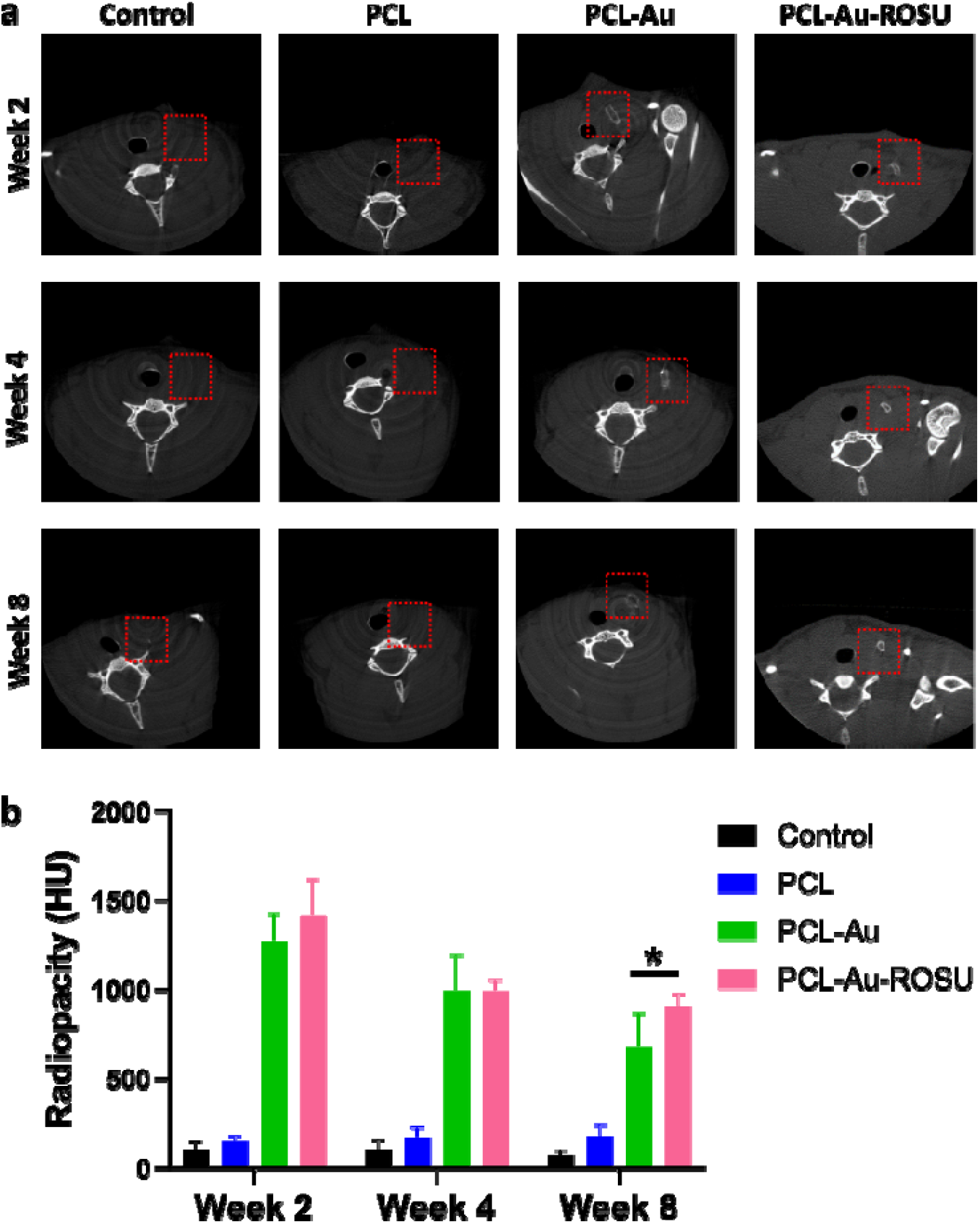
Micro-Computed Tomographic Analysis of Perivascular Wraps. (a) In vivo mCT images of the AVF groups (red box=AVF location) and (b) HU values of the wraps at weeks 2, 4, and 8. AuNP-loading confers enhanced visibility of wraps on mCT and quantitatively greater radiodensity values compared to bare PCL and control at all time points. AuNP-associated radiopacity decreased over time as the in vivo wraps degraded. The addition of ROSU to AuNP-loaded wraps did not reduce wrap visibility, and at week 8, PCL-Au-ROSU was found to be significantly more radiopaque than PCL-Au alone. A significant difference (p<0.05) is indicated by a (*)significance between AuNP wraps versus control, and PCL wraps not annotated for figure readability. Abbreviations: Au, gold; PCL, polycaprolactone, ROSU, rosuvastatin; mCT, micro-computed tomography; HU, Hounsfield units,

### ROSU loading of perivascular wraps demonstrates reduced venous NIH on US analysis

Representative US images of the AVF site across each treatment group and time point are presented in figure 6(A). Average WT:LD ratios, included in supplementary table 3 and figure 6(b), found that this ratio - a marker of NIH - decreased over time in all conditions with a perivascular wrap and increased over time in the control. Wraps with ROSU were found to be equivalent to other perivascular wrap treatment groups at W4 and, notably, had a significantly lower WT:LD ratio than all other cohorts by W8. At W4, the average WT:LD ratio of the PCL-Au-ROSU group was reduced compared to the control (p<0.0001) and was equivalent to the other wrap groups, PCL and PCL-Au (p≥0.4575). By W8, the nanoparticle-and-drug loaded PCL-Au-ROSU group had significantly less NIH by WT:LD ratio than the control (p< 0.0001 and PCL and PCL-Au (p<0.0001).

**Figure 6.**
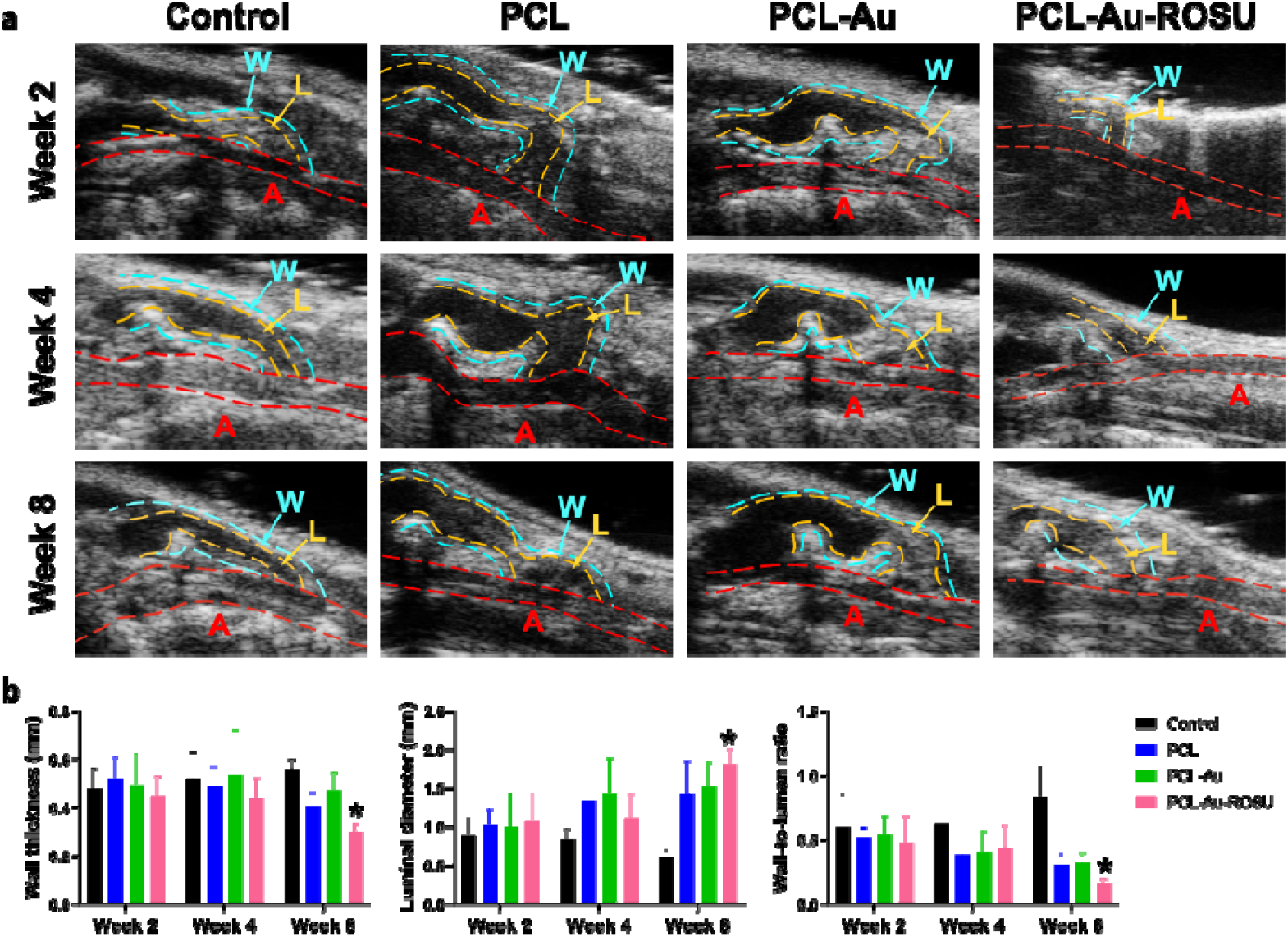
Ultrasound Analysis of Perivascular Wraps. (a) B-mode US images of the AVF groups (A, artery, L, lumen of vein, W, wall of vein, bar=1 mm), and (b) B-mode measurements of the AVF groups at weeks 2, 4, and 8. Decreasing LD, increasing WT, and increasing WT:LD ratio is markers of NIH and loss of patency. As seen in the representative images, venous outflow track lumens increase in size over time in implanted wrap groups but show narrowing by week 8 in control. The quantified analysis further demonstrated this trend and decreased wall thickness and WT:LD ratio in wrap groups. By week 4, all implanted wrap groups showed significantly reduced markers of NIH and occlusion compared to the control. By week 8, ROSU-loaded wraps had significantly increased vessel patency when compared to wraps without ROSU loading [statistical significance (p<0.05) indicated by a (*)]. Abbreviations: Au, gold; PCL, polycaprolactone, ROSU, rosuvastatin; A, artery; V, vein; W, wall;

No differences in WT:LD ratio was seen between any groups at W2 (p≥0.0715), but all groups with implanted perivascular wraps - PCL, PCL-Au, and PCL-Au-ROSU - demonstrated significantly reduced NIA:LA ratios compared to the control by W4 (p<0.0001), and again at W8 (p<0.0001). Finally, there was no significant difference between PCL and PCL-Au at any timepoint (p≥0.8908).

### The addition of ROSU to perivascular wraps demonstrates equivalent to enhanced attenuation of NIH on histologic analysis

Figure 7(a) provides representative images of histological slides with analysis tracings across all groups and time points. Similar to the trends seen on US analysis, NIA:LA ratio decreased over time in the perivascular wrap groups and increases in the control group, with average values included in supplementary table 5 and plotted in figure 7(b). As on US, W4 PCL-Au-ROSU demonstrated a significantly lower NIA:LA ratio as compared to control (p=0.0242) and was equivalent to the other perivascular wrap groups, PCL and PCL-Au (p≥0.1471). By W8, PCL-Au-ROSU was found to have a significantly lower NIA:LA ratio than both control (p=0.0150) and the wraps without ROSU addition, PCL and PCL-Au (p≤0.0269).

**Figure 7.**
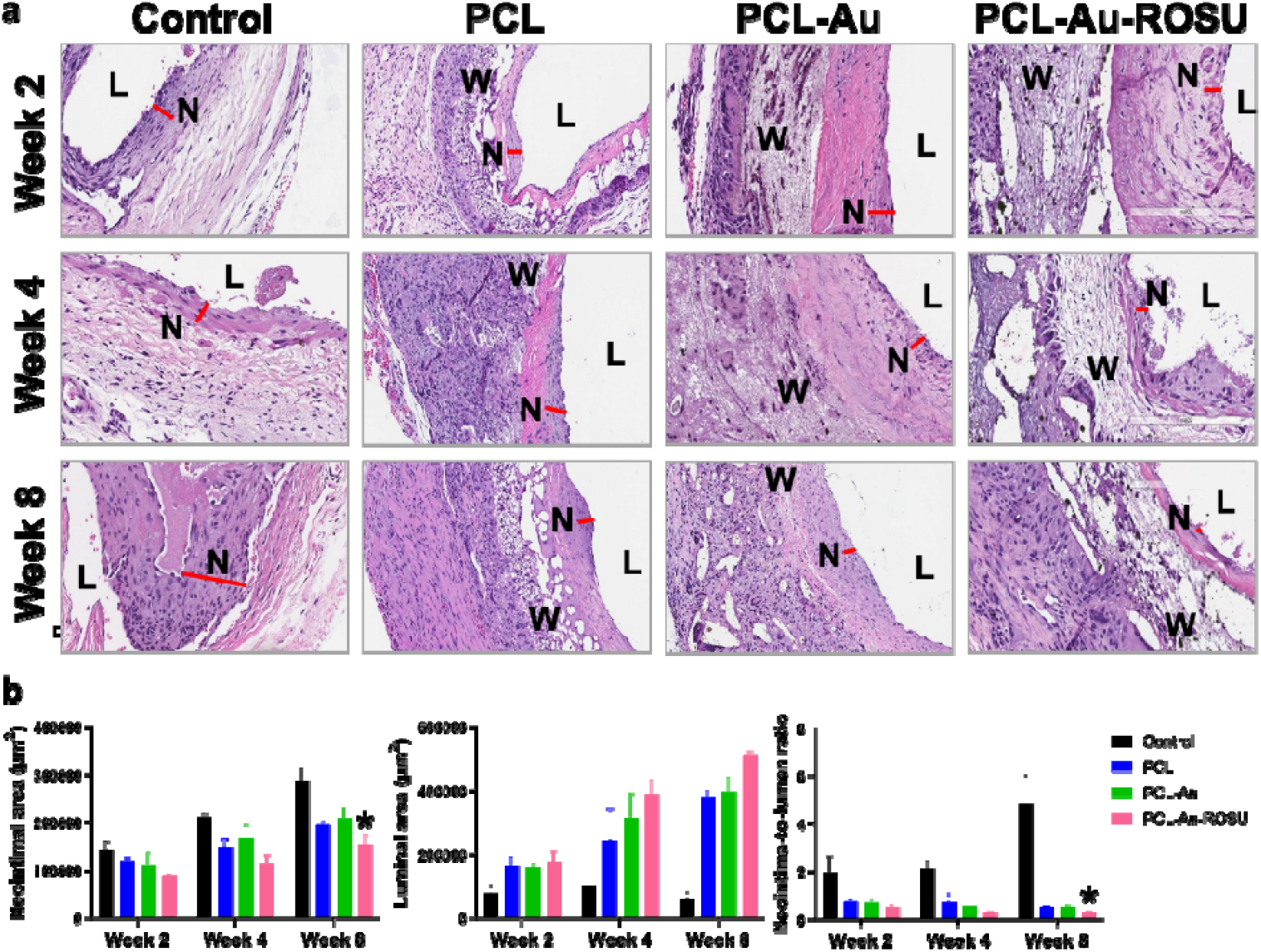
Histological Analysis of Perivascular Wraps. (a) Hematoxylin and Eosin microscopic images of the AVF groups (L, lumen of the vein, N, neointima of the vein, W, wrap, bar=200 um), and (b) histomorphometric measurements of the AVF groups at weeks 2, 4, and 8. The representative histologic images demonstrate an increasing spread of NIH throughout the vessel well and decreased lumen patency at 8 weeks in the control group AVFs. Groups with implanted wrap were found to have reduced NIH and increasing luminal patency over time. These findings were quantified as NIH area, luminal area, and NIH-to-luminal area ratio, which found that wrap groups demonstrated better vessel preservation by week 4 and ROSU-loaded wraps outperformed wraps without ROSU by week 8 [statistical significance (p<0.05) indicated by a (*)]. Abbreviations: Au, gold; PCL, polycaprolactone, ROSU, rosuvastatin; L, lumen of vein; N, neointima of vein; W, wrap.

Although PCL-Au-ROSU was found to have the greatest NIA:LA attenuation by W8, all perivascular wrap groups demonstrated consistently decreased NIA:LA ratios relative to control by W4 (p≤0.0244) and again at W8 (p≤0.0173). As on US, W2 analysis found no difference in NIA:LA between any cohorts (p≥0.1474), and perivascular wraps without ROSU, PCL, and PCL-Au, were found to have equivalent NIA:LA ratios at all time points (p≥0.8721).

## DISCUSSION

In this study, we have combined two potential modifying factors of AVF outcome into a single intervention—a supportive perivascular wrap loaded with AuNPs for enhanced radiologic visibility and ROSU for greater NIH attenuation and vessel patency. We found that the radiopaque perivascular wrap, like the bare wrap, provided a benefit on vessel lumen and wall integrity after AVF creation, but the addition of ROSU—a lipid-lowering agent involved in reduction of vascular inflammation and thrombogenicity (5)—further enhanced this benefit, outperforming the wraps without drug elution.

Our initial results showed that the addition of AuNPs and ROSU influenced the polymeric wrap’s pore size and mechanical strength. The wraps infused with AuNPs alone had larger average pore size, which is associated with greater rates of particle dispersion (19,20). However, there was no difference found in wrap porosity by intrusion method or *in vitro* AuNP release, suggesting that the difference in pore diameter did not impact the rate of substrate intrusion. Though average wrap mechanical strength did decrease with the addition of both AuNPs and ROSU, it remained greater than 5 MPa (15), which is the upper limit of average tensile strength of naturally occurring blood vessels. Therefore, all wrap groups were found to have the sufficient mechanical strength to support vascular flow. In addition, the increased fiber alignment seen in wraps loaded with both AuNPs and ROSU is an optimal structure for directing tissue growth, as the micro-patterning associated with alignment more effectively reconstitutes the form and function found in a native tissue environment (21).

We, then, found that the incorporation of AuNPs significantly increased wrap radiopacity on *in vivo* mCT at all time points, which supports the hypothesis that AuNP-loading confers greater radiographic visibility. Early radiodensity values of the PCL-Au (1,232 ± 75 HU) and PCL-Au-ROSU (1,182 ± 203 HU) wraps are concordant with previous analyses of AuNP radiodensities exceeding 1,000 HU (22,23). Wraps with AuNPs were not only more visible but also demonstrated linear decreases in both *in vitro* AuNP release and *in vivo* radiopacity from W1 to W4. These trends raise the possibility of a predictive model for wrap degradation based on non-invasive clinical imaging. The addition of ROSU to AuNP-loaded wraps showed no decrease in visibility and was found to increase wrap radiopacity at W8, likely due to the ROSU salt-containing calcium, which has similar radiodensity to bone - around 1,000 HU (24). *In vitro* analysis showed that ROSU elution follows a typical pharmacokinetic pattern of release, with an initial burst that tapers off for sustained release over 12 weeks. The greater cumulative loss of AuNPs (88.2-92.4%) compared to ROSU (40.7%) seen in the release studies projects a widening proportion of retained ROSU to retained AuNPs over time, potentially causing the delayed variation in radiopacity.

Lastly, US and histologic analyses revealed a potential therapeutic synergism between the supportive perivascular wrap and ROSU. All the perivascular wraps were found to have a significant advantage by W4, with improved vessel patency and wall structure in all wrap groups. This is consistent with previous reports that perivascular wraps confer a mechanical effect on the vessel wall that suppresses abnormal cell proliferation and vascular changes (8).The addition of ROSU further enhanced the vascular benefits of perivascular wraps by W8, as wraps with ROSU had significantly decreased markers of NIH and luminal narrowing on analysis of primary outcome ratios and isolated subcomponents. This may be explained by the ROSU-loaded wrap’s two-pronged approach to reducing NIH. The mechanical mechanism of NIH attenuation that is inherently present in ROSU-loaded wraps may be bolstered by the additional biochemical mechanism of direct statin delivery to reduce low-density lipoprotein levels, decrease inflammation, and mitigate endothelial cell dysfunction in local tissues. By targeting abnormal vessel changes through two unique pathways, the ROSU-loaded wraps demonstrated improved luminal maintenance and wall integrity than wraps that solely provide mechanical benefit. These results suggest that targeted delivery of ROSU can supplement the NIH-attenuating effect of perivascular wraps, leading to even greater control of vessel patency and, therefore, maintenance of fistula access. Nonetheless, the scope of this study is limited to evaluating the effect of the wraps on NIH as seen in ultrasonography and histomorphometry, and further studies are needed to elucidate the exact mechanisms that underlie the efficacy of these treatments.

## CONCLUSION

Our findings support the increased radiologic visibility of AVF wraps with AuNP loading and demonstrate no loss of radiopacity on mCT with the addition of ROSU to AuNP-loaded wraps. AuNP release and the resulting decrease in radiopacity were both found to follow a linear trend for 4 weeks, allowing for more accurate predictivity of wrap degradation through serial imaging. Additionally, ROSU loading significantly increased venous outflow diameter and further reduced NIH on US and histological analysis at W8 compared to both control and wraps without ROSU loading. These findings suggest that adding ROSU to implanted perivascular wraps for targeted elution may have a significant therapeutic benefit in attenuating post-AVF creation vascular changes and maintaining fistula patency for long-term access.

### Clinical Perspectives

### Competency in Medical Knowledge

Vessel stenosis driven by neointimal hyperplasia is one of the leading complications affecting arteriovenous fistula maturation in patients requiring hemodialysis.

Competency in Patient Care and Procedural Skills: N/A

Competency in Interpersonal and Communication Skills: N/A

Competency in Systems-Based Practice: N/A

Competency in Practice-Based Learning: N/A

Competency in Professionalism: N/A

### Translational Outlook

**Translational Outlook 1:** The structural functionality of perivascular wraps used to supplement arteriovenous fistula physiology is enhanced by incorporating additional, localized pharmacologic therapy such as rosuvastatin.

**Translational Outlook 2:** The therapeutic mechanical and pharmacological effects of bioabsorbable perivascular polycaprolactone wraps are more effectively monitored using non-invasive imaging modalities when supplemented with radiopaque gold nanoparticles.

## Abbreviations

AVF: Arteriovenous Fistula
NIH: Neointimal Hyperplasia
ROSU: Rosuvastatin
AuNP: Gold Nanoparticle
PCL: Polycaprolactone
mCT: Micro-Computed Tomography
HU: Hounsfield Units
W: Week
US: Ultrasound
WT: Wall Thickness
LD: Luminal Diameter
LA: Luminal Area
NIA: Neointimal Area

## Acknowledgments

The authors would like to acknowledge Sunita Paterson in MD Anderson’s Research Medical Library for editing the manuscript, Dr. James Gu at the Electron Microscopy Core at Houston Methodist Research Institute for assisting with the conduct of scanning electron microscopy, Dunn Lab personnel (i.e., Amanda McWatters, Malea L. Williams, and Steve D. Parrish) for assisting with animal experiments, and Small Animal Imaging Facility personnel for assisting with animal imaging.

## Figure Titles and Legends (including Central Illustration)

## Supplementary Material

**Supplementary Table 1.** In vitro (a) % drug release and (b) % AuNP release *Abbreviations: Au, gold; PCL, polycaprolactone, ROSU, rosuvastatin*

| Average % ROSU Release |  |  |  |
| --- | --- | --- | --- |
| Week | PCL | PCL-Au | PCL-Au-ROSU |
| 0 | $0 \pm 0$ | $0 \pm 0$ | $0 \pm 0$ |
| 1 | $0 \pm 0$ | $0 \pm 0$ | $26.5 \pm 16.9$ |
| 2 | $0 \pm 0$ | $0 \pm 0$ | $29.2 \pm 14.9$ |
| 3 | $0 \pm 0$ | $0 \pm 0$ | $30.9 \pm 13.5$ |
| 4 | $0 \pm 0$ | $0 \pm 0$ | $32.2 \pm 13.2$ |
| 5 | $0 \pm 0$ | $0 \pm 0$ | $33.1 \pm 13.2$ |
| 6 | $0 \pm 0$ | $0 \pm 0$ | $33.9 \pm 13.2$ |
| 8 | $0 \pm 0$ | $0 \pm 0$ | $37.9 \pm 12.4$ |
| 10 | $0 \pm 0$ | $0 \pm 0$ | $39.4 \pm 12.7$ |
| 12 | $0 \pm 0$ | $0 \pm 0$ | $40.7 \pm 13.0$ |

**Supplementary Table 2.** In vitro % AuNP release *Abbreviations: Au, gold; AuNP, gold nanoparticle; PCL, polycaprolactone, ROSU, rosuvastatin*

| Average % AuNP Release |  |  |  |
| --- | --- | --- | --- |
| Week | PCL | PCL-Au | PCL-Au-ROSU |
| 0 | $0 \pm 0$ | $0 \pm 0$ | $0 \pm 0$ |
| 1 | $0 \pm 0$ | $18.76 \pm 0.16$ | $19.384 \pm 0.69$ |
| **2** | $0 \pm 0$ | $28.36 \pm 0.24$ | $29.25 \pm 1.04$ |
| **3** | $0 \pm 0$ | $47.39 \pm 0.40$ | $47.186 \pm 1.42$ |
| **4** | $0 \pm 0$ | $58.56 \pm 0.40$ | $57.875 \pm 0.36$ |
| **8** | $0 \pm 0$ | $73.34 \pm 0.44$ | $77.604 \pm 5.16$ |
| **12** | $0 \pm 0$ | $88.18 \pm 0.55$ | $92.42 \pm 5.22$ |

**Supplementary Table 3.** In vitro (a) RF24 and (b) MOVAS cell line viability *Abbreviations: Au, gold; PCL, polycaprolactone, ROSU, rosuvastatin*

| **RF24 Cell Viability (%)** |  |  |  |  |
| --- | --- | --- | --- | --- |
| **Treatment (%)** | **Control** | **PCL** | **PCL-Au** | **PCL-Au-ROSU** |
| **25** | $1 \pm 0.351$ | $1.074 \pm 0.081$ | $0.928 \pm 0.03$ | $0.911 \pm 0.053$ |
| **50** | $0.999 \pm 0.142$ | $1.147 \pm 0.146$ | $0.934 \pm 0.123$ | $0.77 \pm 0.159$ |
| **75** | $1 \pm 0.084$ | $1.107 \pm 0.433$ | $0.829 \pm 0.15$ | $0.807 \pm 0.105$ |
| **100** | $0.999 \pm 0.18$ | $0.845 \pm 0.288$ | $1.096 \pm 0.298$ | $0.898 \pm 0.373$ |
| **MOVAS Cell Viability (%)** |  |  |  |  |
| **Treatment (%)** | **Control (N=3)** | **PCL** | **PCL-Au** | **PCL-Au-ROSU** |
| **25** | $0.999 \pm 0.104$ | $1.095 \pm 0.06$ | $1.007 \pm 0.287$ | $0.897 \pm 0.115$ |
| **50** | $1 \pm 0.394$ | $0.874 \pm 0.138$ | $0.97 \pm 0.197$ | $0.927 \pm 0.052$ |
| **75** | $1 \pm 0.15$ | $1.319 \pm 0.389$ | $1 \pm 0.147$ | $1.194 \pm 0.165$ |
| **100** | $1 \pm 0.195$ | $1.055 \pm 0.235$ | $1.044 \pm 0.35$ | $0.999 \pm 0.195$ |

**Supplementary Table 4.** In vivo mCT radiopacity of AVF groups *Abbreviations: Au, gold; HU, Hounsfield unit; PCL, polycaprolactone, ROSU, rosuvastatin*

| Average Radiopacity (HU) |  |  |  |  |
| --- | --- | --- | --- | --- |
| Week | Control | PCL | PCL-Au | PCL-Au-ROSU |
| 2 | 78.1 ± 32.3 | 143.5 ± 22.4 | 1232.3 ± 75.7 | 1182.0 ± 203.9 |
| 4 | 75.3 ± 21.3 | 61.1 ± 14.9 | 939.1 ± 166.4 | 1006.8 ± 57.8 |
| 8 | 73.7 ± 27.7 | 61.2 ± 29.9 | 573.8 ± 132.3 | 895.4 ± 77.8 |

**Supplementary Table 5.**
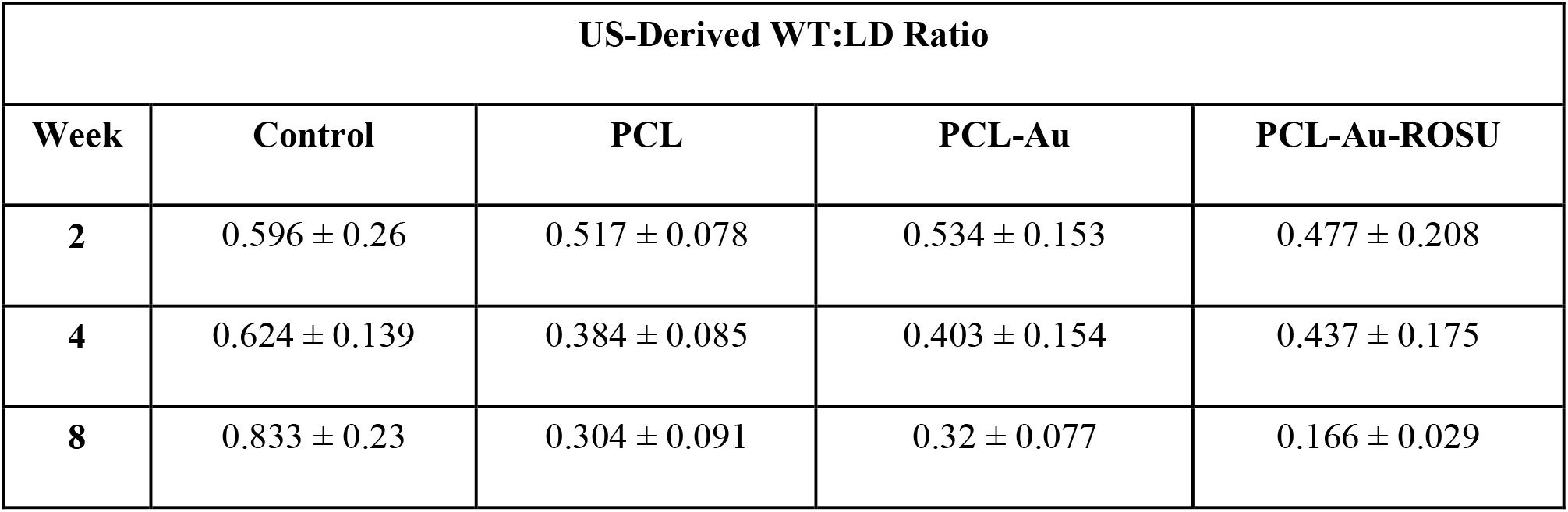
In vivo US-derived wall-to-lumen ratios of AVF groups *Abbreviations: Au, gold; LD, luminal diameter; PCL, polycaprolactone, ROSU, rosuvastatin; WT, wall thickness*

**Supplementary Table 6.** In vivo histologic NIH-to-lumen ratios of AVF groups *Abbreviations: Au, gold; LA, luminal area; NIA, neointimal area; PCL, polycaprolactone, ROSU, rosuvastatin*

| Histologic NIA:LA Ratio |  |  |  |  |
| --- | --- | --- | --- | --- |
| Week | Control | PCL | PCL-Au | PCL-Au-ROSU |
| 2 | 1.97 ± 0.667 | 0.737 ± 0.109 | 0.685 ± 0.134 | 0.51 ± 0.084 |
| 4 | 2.116 ± 0.315 | 0.706 ± 0.36 | 0.543 ± 0.113 | 0.293 ± 0.015 |
| 8 | 4.84 ± 1.216 | 0.52 ± 0.014 | 0.53 ± 0.06 | 0.3 ± 0.034 |

**Online Figure 1.**
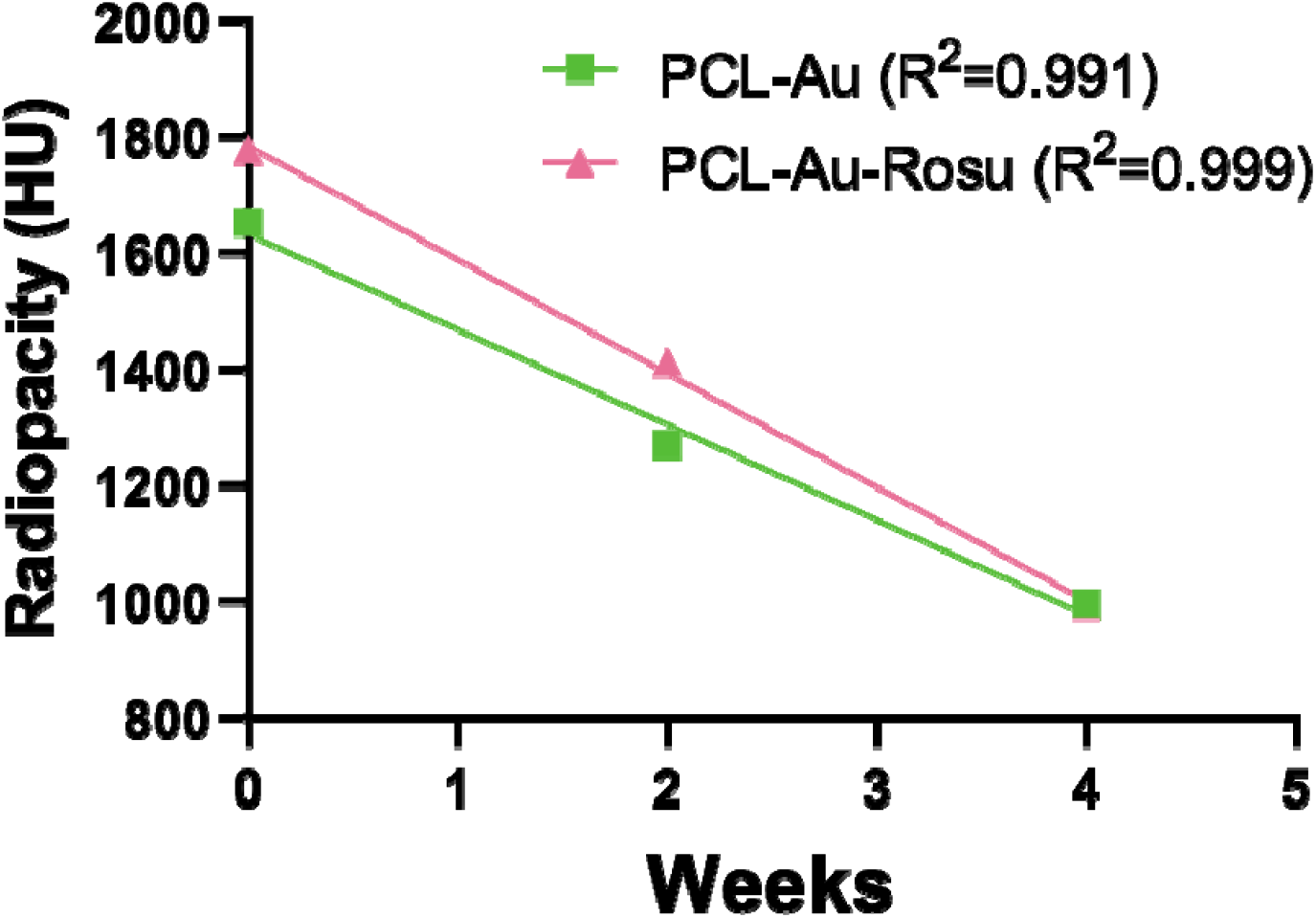
*In vivo* HU values of AuNP-containing wraps up to 4 weeks. Abbreviations: Au, gold; PCL, polycaprolactone, ROSU, rosuvastatin

## Notes

***Funding*** This research was funded by the National Institutes of Health – National Heart, Lung, and Blood Institute (5HL141831-05 and 1R01HL159960-01A1), a Radiological Society of North America Research Seed Grant (RSD2012), a Society of Interventional Radiology Pilot Research Grant, MD Anderson’s Center for Advanced Biomedical Imaging Pilot Project Program Research Grant, and the National Institutes of Health – National Cancer Institute through MD Anderson’s Cancer Center Support Grant (P30CA016672; used the Research Animal Support Facility and Small Animal Imaging Facility).

### Competing Interest Statement

The authors have declared no competing interest.

## References

1. United States Renal Data System. 2022 USRDS Annual Data Report: Epidemiology of kidney disease in the United States. National Institutes of Health, National Institute of Diabetes and Digestive and Kidney Diseases, Bethesda, MD, 2022.

2. Cheung AK, Imrey PB, Alpers CE, et al. Intimal Hyperplasia, Stenosis, and Arteriovenous Fistula Maturation Failure in the Hemodialysis Fistula Maturation Study. Journal of the American Society of Nephrology. 2017;28(10):3005–3013.

3. Ma S, Duan S, Liu Y, Wang H. Intimal Hyperplasia of Arteriovenous Fistula. Annals of Vascular Surgery. 2022;85:444–453.

4. Yamazaki T, Shirai H, Yashima J, Tojimbara T. High low-density lipoprotein cholesterol level is the independent risk factor of primary patency rate of arteriovenous fistula. Vascular. 2020;28(4):430–435.

5. Chang H-H, Chang Y-K, Lu C-W, et al. Statins Improve Long Term Patency of Arteriovenous Fistula for Hemodialysis. Scientific Reports. 2016;6(1):22197.

6. Wan Q, Li L, Yang S, Chu F. Impact of Statins on Arteriovenous Fistulas Outcomes: A Meta-Analysis. Therapeutic Apheresis and Dialysis. 2018;22(1):67–72.

7. Martinez L, Duque JC, Escobar LA, et al. Distinct Impact of Three Different Statins on Arteriovenous Fistula Outcomes: A Retrospective Analysis. The Journal of Vascular Access. 2016;17(6):471–476.

8. Barcena AJR, Perez JVD, Liu O, et al. Localized Perivascular Therapeutic Approaches to Inhibit Venous Neointimal Hyperplasia in Arteriovenous Fistula Access for Hemodialysis Use. Biomolecules. 2022;12(10):1367.

9. San Valentin EMD, Barcena AJR, Klusman C, Martin B, Melancon MP. Nano-embedded medical devices and delivery systems in interventional radiology. Wiley Interdisciplinary Reviews Nanomedicine and Nanobiotechnology. 2023;15(1):e1841.

10. Goldstein A, Soroka Y, Frušić-Zlotkin M, Popov I, Kohen R. High resolution SEM imaging of gold nanoparticles in cells and tissues. Journal of Microscopy. 2014;256(3):237–247.

11. Damasco JA, Huang SY, Perez JVD, et al. Bismuth Nanoparticle and Polyhydroxybutyrate Coatings Enhance the Radiopacity of Absorbable Inferior Vena Cava Filters for Fluoroscopy-Guided Placement and Longitudinal Computed Tomography Monitoring in Pigs. ACS Biomaterials Science & Engineering. 2022;8(4):1676–1685.

12. Huang SY, Damasco JA, Tian L, et al. In vivo performance of gold nanoparticle-loaded absorbable inferior vena cava filters in a swine model. Biomaterials Science. 2020;8(14):3966–3978.

13. Tian L, Lee P, Singhana B, et al. In vivo imaging of radiopaque resorbable inferior vena cava filter infused with gold nanoparticles. Proceedings of SPIE--the International Society for Optical Engineering. 2018;10576:105762S.

14. Tian L, Lee P, Singhana B, et al. Radiopaque Resorbable Inferior Vena Cava Filter Infused with Gold Nanoparticles. Scientific reports. 2017;7(1):2147–2110.

15. Zhu X, Cui W, Li X, Jin Y. Electrospun Fibrous Mats with High Porosity as Potential Scaffolds for Skin Tissue Engineering. Biomacromolecules. 2008;9(7):1795–1801.

16. Soliman S, Pagliari S, Rinaldi A, et al. Multiscale three-dimensional scaffolds for soft tissue engineering via multimodal electrospinning. Acta biomaterialia. 2010;6(4):1227–1237.

17. Wang X, Chaudhry MA, Nie Y, et al. A Mouse 5/6th Nephrectomy Model That Induces Experimental Uremic Cardiomyopathy. Journal of Visualized Experiments. 2017(129).

18. Wong CY, de Vries MR, Wang Y, et al. A Novel Murine Model of Arteriovenous Fistula Failure: The Surgical Procedure in Detail. Journal of Visualized Experiments. 2016(108):e53294–e53294.

19. Klose D, Siepmann F, Elkharraz K, Krenzlin S, Siepmann J. How porosity and size affect the drug release mechanisms from PLGA-based microparticles. International Journal of Pharmaceutics. 2006;314(2):198–206.

20. Siepmann J, Faisant N, Akiki J, Richard J, Benoit JP. Effect of the size of biodegradable microparticles on drug release: experiment and theory. Journal of Controlled Release. 2004;96(1):123–134.

21. Baker BM, Mauck RL. The Effect of Nanofiber Alignment on the Maturation of Engineered Meniscus Constructs. Biomaterials. 2007;28(11):1967–1977.

22. Mesbahi A, Famouri F, Ahar MJ, Ghaffari MO, Ghavami SM. A study on the imaging characteristics of Gold nanoparticles as a contrast agent in X-ray computed tomography. Polish Journal of Medical Physics and Engineering. 2017;23(1):9–14.

23. Silvestri A, Zambelli V, Ferretti AM, et al. Design of functionalized gold nanoparticle probes for computed tomography imaging. Contrast Media & Molecular Imaging. 2016;11(5):405–414.

24. Lev MH, Gonzalez RG. 17 - CT Angiography and CT Perfusion Imaging. In: Toga AW, Mazziotta JC, eds. Brain Mapping: The Methods (Second Edition). San Diego: Academic Press; 2002:427–484.

